# The Role of Locus Coeruleus and Substantia Nigra/Ventral Tegmental Area activity in Reward and Salience Processing: A High-Resolution fMRI Study

**DOI:** 10.64898/2026.10.08.752674

**Authors:** J. M. Hall, N. Kukkonen, E. Prinea, J. Schomaker, R. M. Krebs

**Affiliations:** Department of Health, Medical and Neuropsychology, Leiden University, Leiden, the Netherlands; Department of General and Applied Psychology, Ghent University, Ghent, Belgium; Leiden University Medical Centre, Leiden, The Netherlands

**Keywords:** reward, saliency, dopamine, noradrenaline, fMRI, oddball

## Abstract

Dopaminergic neurons in the substantia nigra/ventral tegmental area (SN/VTA) are classically associated with reward processing, whereas noradrenergic neurons in the locus coeruleus (LC) have been shown to support salience processing more broadly. However, rewarding stimuli are also inherently salient, and in most paradigms reward and salience are conflated, for example when rewarded targets are also rare and/or visually distinct. As a result, it remains unclear in how far these systems are sensitive to reward-specific properties or more basic salient properties of the stimulus. Using a visual oddball paradigm with high-resolution fMRI (n = 45), we compared oddballs that were rewarded, visually salient, or neutral, while keeping stimulus probability and motor requirements constant across conditions. Participants responded more accurately to rewarded than to neutral oddballs, and faster to rewarded than to both neutral and visually salient oddballs. At the neural level, SN/VTA activity was increased for rewarded oddballs compared with both neutral and visually salient oddballs, whereas LC activity was increased for rewarded compared with visually salient oddballs. Visual salience alone did not increase activity in either region relative to neutral oddballs. Together, these findings suggest that both the noradrenergic and dopaminergic systems are preferentially engaged by motivationally relevant stimuli, beyond more basic forms of salience.

## Introduction

The dopaminergic (DA) system has traditionally been implicated in reward processing, whereas the noradrenergic (NA) system has been most prominently associated with processing salient information (e.g., Sara & Bouret, 2012; Schultz, 2002). However, more recent studies in both animals and humans show substantial overlap between the two systems: NA activity is modulated by reward, and DA activity by salient, non-rewarding events (e.g., Bromberg-Martin et al., 2010; Cole et al., 2022; Varazzani et al., 2015). This raises the question of whether the two systems encode a shared salience signal, or whether they differ in the type of salience to which they respond.

The locus coeruleus (LC) is the primary source of the catecholamine neuromodulator noradrenaline (NA; e.g. Poe et al., 2020). The LC is a small nucleic structure residing in the pons and has widespread connections throughout the brain. It is therefore well positioned to modulate a wide range of different functions depending on its firing rate, modular organisation and target characteristics (Agster et al., 2013; Berridge & Waterhouse, 2003; Grimm et al., 2024; Poe et al., 2020; Sara & Bouret, 2012). One of the key functions of the LC-NA is modulating arousal, attention and effort, that serve goal-directed behaviour. It detects unfamiliar and salient stimuli and orients attention towards them (Maness et al., 2022; Poe et al., 2020; Sara & Bouret, 2012). Salience may arise from physical stimulus characteristics (visual salience), from unexpectedness (e.g., a rare oddball), or from motivational relevance (e.g., association with reward). This initial orienting response allows rapid learning about a stimulus; as the stimulus becomes familiar, the attentional engagement of the LC-NA system decreases. This shift from broad orienting to selective disengagement is thought to reflect a more general property of the LC-NA system: rather than responding to any single stimulus feature, it continuously recalibrates attentional resources according to how informative or relevant a stimulus currently is. This flexibility becomes especially apparent when a stimulus is motivationally relevant (Cole et al., 2022; Nieuwenhuis et al., 2005).

Indeed, both animal (Bouret & Richmond, 2015; Jahn et al., 2018; Varazzani et al., 2015) and human studies (Cole et al., 2022; Hall et al., 2024) have shown the involvement of the LC-NA system in reward processing. In animal models, the LC has been shown to modulate arousal and attention during reward-related tasks, facilitating a rapid adaptation to changing reward stimuli (Bouret & Sara, 2005; Sara & Bouret, 2012). This activation is critical for directing attention and arousal to behaviourally relevant stimuli, thereby ensuring an optimal state for the brain to process information (Sara & Bouret, 2012). But beyond attentional modulation, the LC also contributes to executing reward-related behaviours by scaling the effort allocated to actions in proportion to their expected reward (Bouret & Richmond, 2015; Jahn et al., 2018; Varazzani et al., 2015). For example, LC activity has been linked to adjustments in response vigour and physiological arousal (e.g., pupil dilation) that track the cost-benefit trade-off of pursuing a reward, effectively translating motivational value into the energisation of goal-directed action (Bouret & Richmond, 2015).

Consistent with this role in translating motivational value into goal-directed behaviour, recent human work shows that LC responses are modulated by reward, a function traditionally ascribed to the dopaminergic system. Much of this work has used pupil size as an index of LC activity: the LC influences pupil diameter through its projections to autonomic pathways (Samuels & Szabadi, 2008). Using this measure, Cole et al. (2022) showed that reward incentives amplified both tonic baseline pupil size and phasic pupil responses to oddball stimuli (low-frequency target stimuli that require a response) in high-reward compared with low-reward contexts. This is consistent with enhanced LC engagement beyond stimulus salience alone. Beyond reward, however, emotional valence is also motivationally relevant, as emotional stimuli capture attention and engage approach or avoidance tendencies (Lang & Bradley, 2010). Human high-resolution fMRI work further suggests that LC responses are sensitive to the valence and task-relevance of events: LC activation was higher for loss than for gain feedback, and for feedback signalling a task reversal, particularly in older adults (Ludwig et al., 2024). Extending these findings, we recently investigated LC-NA involvement in an fMRI reward-valence recognition task (Hall et al., 2024). In this task, participants needed to respond as accurately and quickly as possible to identify the emotional expression of a face (happy, neutral or angry), to earn a reward. Reward availability was signalled independently by the gender of the face, while emotional valence was signalled by its expression. This allowed us to dissociate reward anticipation from emotional valence within the same stimuli. We confirmed that LC activity increased during reward-anticipatory trials in which a behavioural response was required. Notably, this activity was further amplified for reward trials with additional positive valence (i.e., here conveyed via happy facial expressions) revealing an additive effect of reward and positive valence on LC activation.

Reward and positive-valence-related processing has traditionally been linked to the dopaminergic substantia nigra/ventral tegmental area complex (SN/VTA). Dopamine facilitates reinforcement learning by signalling prediction errors, such that learning occurs when outcomes deviate from expectations (Schultz, 2002). Furthermore, DA is involved in processing rewards and motivating behaviour into goal-directed actions (Schultz, 2002, 2013; Schultz et al., 1997; Wise, 2004) including the vigour with which those actions are performed (Hofmans et al., 2022). This is accomplished by enhancing the incentive salience of reward-related stimuli, and thus influencing the individual’s drive to pursue those rewards, ultimately playing a crucial role in motivation (Berridge, 2008).

In our recent study (Hall et al., 2024), the SN/VTA also responded to salient stimuli when a response was required. Activity increased on rewarded trials with neutral valence, but also on non-rewarded trials with negative valence (compared to neutral). This is in line with animal work showing that DA neurons respond to aversive events (Brischoux et al., 2009; Matsumoto & Hikosaka, 2009). Such responses are thought to reflect motivational salience, which is signalled for both rewarding and aversive events, as well as alerting signals to unexpected events (Bromberg-Martin et al., 2010). Thus, both LC-NA and SN/VTA-DA systems were sensitive to reward, with distinct valence response profiles across the two neuromodulatory systems.

A key open question is therefore whether LC and SN/VTA responses reflect reward itself or the salience that accompanies it. In previous paradigms, these two have been confounded. In Hall et al. (2024), reward and valence were equally task-relevant, so both conveyed salience. The same problem arises in rewarded oddball designs. In Cole et al. (2022), reward enhanced pupil responses to oddballs, but these oddballs were both rare and task-relevant targets. The reward related enhancement could therefore reflect perceptual (auditory) salience, motivational salience, or both. Both systems are thought to be influenced by overall stimulus salience (Pessoa, 2009), so separating these sources is necessary to determine whether the NA and DA systems play overlapping or distinct roles.

To systematically investigate the contribution of the NA and DA system in processing reward-related and visual salience (i.e., not related to reward or valence), we employed a visual oddball design while assessing neural activity from LC and SN/VTA with high-resolution fMRI. This design allowed us to directly compare responses to salience in the absence of reward with responses to reward in the absence of oddball salience and compare LC and SN/VTA responses to visual salience versus rewarding stimuli and test whether these neuromodulatory systems play distinct roles in processing salience and reward.

### Methods

#### 2.1.1 Participants

Our power analysis (see https://osf.io/2rsye) indicated the need for 44 participants in the current study. To account for exclusions and drop-out, 53 participants between the ages of 18 and 35 years were recruited through the Ghent University online recruiting website and social media websites. Eight participants were excluded from the imaging analysis due to incomplete scanning sessions or technical issues. Two participants were excluded from the behavioural analyses due to a technical issue during data acquisition, resulting in a final behavioural sample of n = 43. The final imaging sample consisted of 45 participants with a mean age of 23.45 years (SD=3.46y; 62% females). The inclusion criteria included (in addition to the safety screening for MRI participation), right-handedness, normal/corrected vision, and no previous diagnosis of psychiatric disorders. Participation lasted approximately 1.5 hours, after which participants received 30 euros as compensation, with the opportunity to earn an additional 10 euros based on their performance on the reward task. Before the study commenced, all participants provided written informed consent. Ethical approval was obtained from Ghent University Hospital on 25/01/2022 (Ethics approval number: EC/2019/0956).

#### 2.1.2 Experimental design

Participants performed a reward-oddball task in the fMRI scanner. Different geometric shapes were presented to participants, representing a rewarded oddball, a neutral oddball, a visually salient oddball, and a standard stimulus which was nine times more frequent than the other stimuli (Figure 1). The standard condition entailed the presentation of a grey diamond, the rewarded oddball was a grey triangle, the neutral oddball was a flipped grey triangle, and the visually salient oddball was a dark yellow triangle in the same orientation as the neutral oddball. The mapping between condition and triangle orientation was counterbalanced across participants, such that each orientation served equally often as the rewarded, and neutral and visual salient oddball across the sample. All shapes were iso-luminant to ameliorate differences in low-level perceptual salience. Participants were instructed to respond to all oddball stimuli by pressing a dedicated button using their index, ring, or middle fingers, the mapping of which was counterbalanced across participants. Each oddball was presented 15 times each (8%), while the standard stimulus was presented 135 times (75%), resulting in a total of 180 shape presentations per experimental block. Each stimulus was presented for 1000 ms. Participants were instructed to quickly respond with a cut off time of 500 ms, however, we considered responses up to 1000 ms as correct. The time interval between the events was distributed based on multiples of the scan repetition time (TR), with 80% at 1 TR (2090 ms), 10% at 2 TRs (4180 ms), 5% at both 3 TRs (6270 ms) and 4 TRs (8360 ms), to allow for effective event-related BOLD response estimation across trials (Hinrichs et al., 2000). A fixation cross was displayed between stimulus presentations. Participants completed three experimental blocks lasting approximately 10 minutes each, which were programmed using PsychoPy version 2.3 (Python version 3.8; Peirce, 2007). Feedback was provided on screen after each block indicating average accuracy across trials. Note that this feedback display was the same for all participants, gradually increasing with each block to mimic practice effect and promote continuous motivation (79% for Block 1, 83% for Block 2, and 93% for Block 3).

**Figure 1.**
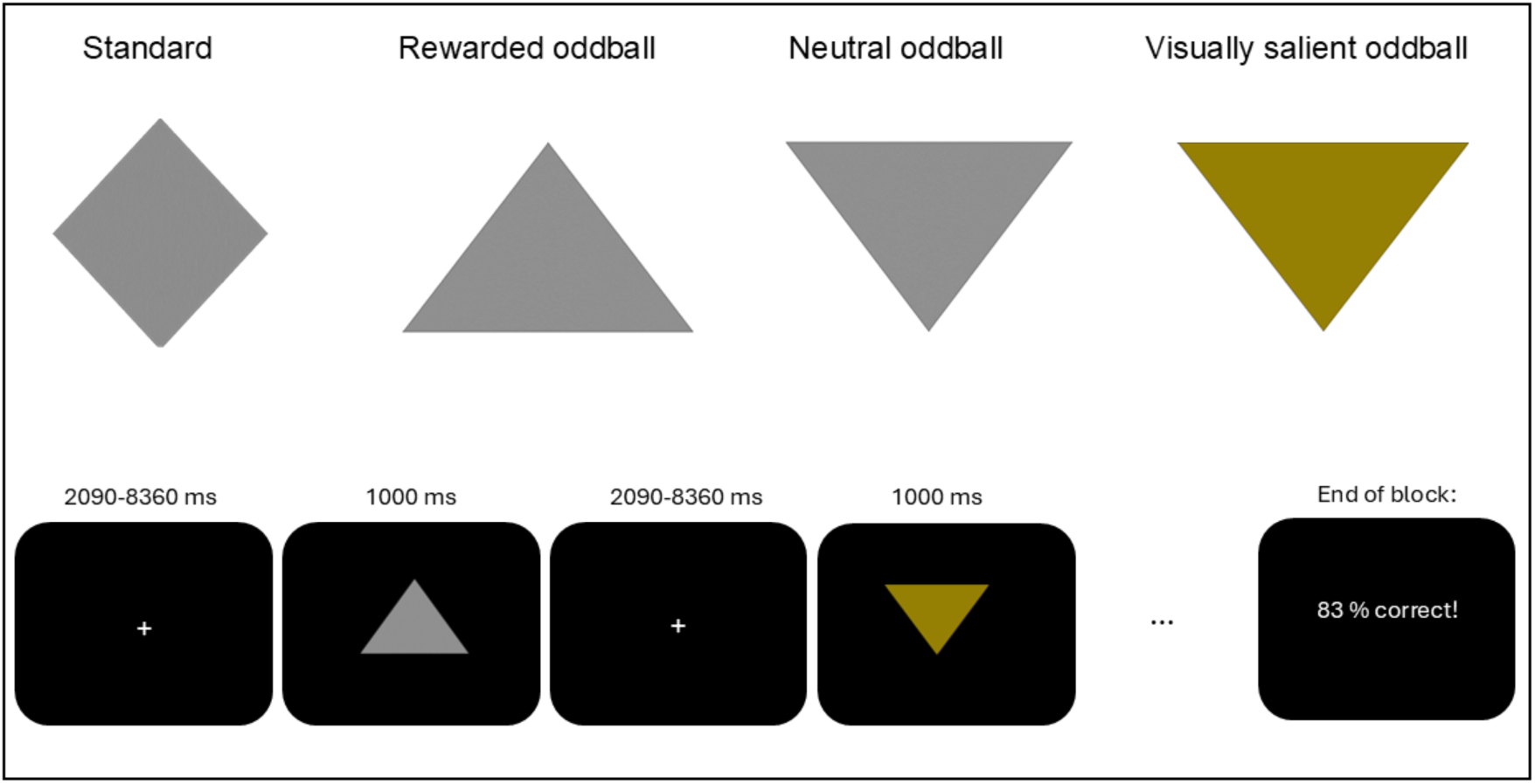
Stimulus types in the reward-oddball task Note. Orientation of rewarded, neutral, and visually salient oddball stimuli were randomised across participants. Only the oddball stimuli (rewarded, neutral and visually salient) required a response.

#### 2.1.3 Image acquisition and pre-processing

MR images were acquired using a 3T Siemens Magnetom Prisma scanner, using a 64-channel head coil. BOLD signal was measured by acquiring T2*-weighted echo-planar functional images in interleaved order with 69 transverse slices (repetition time (TR) = 2090 ms, echo time (TE) = 27 ms, flip angle = 79°, field of view = 112 mm, multiband acceleration factor = 3, no interslice gap, voxel size 1.7 × 1.7 × 2 mm. For the functional localization, a high-resolution T1-weighted structural image was obtained using a 3D magnetisation-prepared rapid acquired gradient echo (MP-RAGE) sequence (TR = 2250 ms, TE = 4.18 ms, FOV = 256 mm, flip angle = 9°; 176 sagittal slices, matrix size = 256 × 256, voxel size = 1 × 1 × 1 mm, duration = 5.14 min). To localize the dopaminergic SN/VTA complex, a T2-weighted structural scan was obtained (TR = 3200 ms; TE = 4.08 ms; FOV = 230 mm; flip angle = 120°; 192 sagittal slices; matrix size = 256 × 256; voxel size = 0.4 × 0.4 × 0.9 mm; duration = 5.09 min). Finally, to localize the LC a neuromelanin-sensitive a T1-weighted turbo spin echo (T1-TSE) scan was acquired (TR = 559 ms; TE = 98 ms; FOV = 192 mm; flip angle 1 = 70°; flip angle 2 = 180°; 10 sagittal slices; matrix size = 384 x 384; voxel size = 0.5 × 0. × 3 mm; duration = 9.51 min).

##### Preprocessing

Images were preprocessed using *fMRIPrep* 21.0.1 (Esteban et al., 2019), which is based on *Nipype* 1.6.1 (Gorgolewski et al., 2011; Gorgolewski et al., 2017). Functional scans were corrected for head motion, slice timing, and co-registered to each participant’s T1-weighted anatomical image. Images were spatially smoothed with a 3.5 mm FWHM Gaussian kernel. Physiological motion confounds, including framewise displacement, DVARS, and CompCor components, were estimated for inclusion in subsequent analyses. Preprocessing of the T1w and T2w scans was performed in fMRIprep 21.0.1, while TSE scans were normalized using Statistical Parametrics Mapping 12 (SPM12; https://www.fil.ion.ucl.ac.uk/spm/software/spm12/). The TSE scans were resliced to a resolution of 0.85*0.85*1.0 mm. The full description of the preprocessing steps is presented in the Supplementary Materials Section A.

#### 2.1.4 Behavioural analyses

The behavioural analyses were performed in Rstudio (R version 4.5.1) using the *afex*and *rstatix* packages. Accuracy data (%) and response times (RT, correct responses only) were averaged across all blocks. For each outcome measure, a repeated-measures analysis of variance (rANOVA) was performed with stimulus type (reward, neutral, visual salience) as the within-subjects factor. Greenhouse-Geisser corrections were applied in cases of sphericity violations and partial eta-squared (η²ₚ) is reported as a measure of effect size. Significant main effects were followed by post hoc paired t-tests, Holm-Bonferroni corrected for multiple comparisons and Cohen’s d is reported as a measure of effect size.

### 2.2 fMRI analysis

#### 2.2.1 First Level Analysis

First-level analysis for each participant was conducted using SPM12. Statistical parametric maps were calculated using a General Linear Model (GLM) with an event-related design. The high-pass filter was set to 128 seconds to remove low-frequency noise and each experimental condition (standard, rewarded, neutral, and visually salient oddball, and errors) was convolved with the canonical haemodynamic response function. Contrast images of participants BOLD response characteristics were obtained for each experimental condition and assigned four, condition specific regressors, and one regressor of noninterest. Six motion regressors were obtained from the realignment procedure as additional regressors of no interest. The focus of this study is on modulations within LC and SN/VTA, however, we have additionally included a whole-brain analysis to provide a complete overview in the Supplementary Materials Section B.

#### 2.2.2 Region of interest analysis

Individual masks were drawn on the structural scans for the SN/VTA (JMH and EP) and LC (JMH and FL) using MRIcroGL (Rorden & Brett, 2000), using the coregistered single subject T2w scans and T1-TSE respectively. Figure 2 shows the localisation of LC and SN/VTA on single-subject anatomical images 2A, and the averaged mask of the nuclei averaged across participants. Our segmentation procedures were carefully guided by anatomical landmarks as well as intensity differences in specific anatomical sequences that were established in previous work on the dopaminergic and noradrenergic systems in the human brain, including our own (Düzel et al., 2009; Hall et al., 2024; Keren et al., 2009; Krebs et al., 2011, 2012, 2018). The LC is visible as bilateral high intensity spots on the axial slices of the T1-TSE scan of the brainstem, just above the fourth ventricle (Fig. 2A, cf. Keren et al., 2009). The SN is visible on the axial slices of the T2w scan as bilateral low intensity (dark) area across the medial midbrain, the VTA lies adjacent to it in anterior-medial direction (Fig. 2A, cf. Düzel et al., 2009). For the purpose of the current study, we only included the ventromedial part of the SN/VTA complex through visual inspection, which has been considered particularly relevant for reward processing (Düzel et al., 2015). Average beta values averaged across each run were extracted using MarsBaR toolbox v0.44 (MARSeille Boîte À Région d’Intérêt; Brett et al., 2002). The individual masks were overlayed with functional scans and converted to an SPM-compatible format. The beta values from each ROI were submitted to an rANOVA with stimulus type (reward, visual salience, neutral) as independent factor. The ‘standard’ stimulus was not included in the main analysis, as no response was required after presentation of this stimulus, hence the results would likely be influenced by these different task requirements. However, an additional rANOVA including the standard was performed as a sanity check. We also used the publicly-available pontine reference area mask as a control region of interest and the beta values of this region was used as a factor in an rANOVA. To assess data quality of the functional signal within the LC ROI, we calculated the temporal signal-to-noise-ratio (tSNR) within the LC by computing the mean value over time, averaged across all voxels of the LC, divided by the standard deviation.

**Figure 2.**
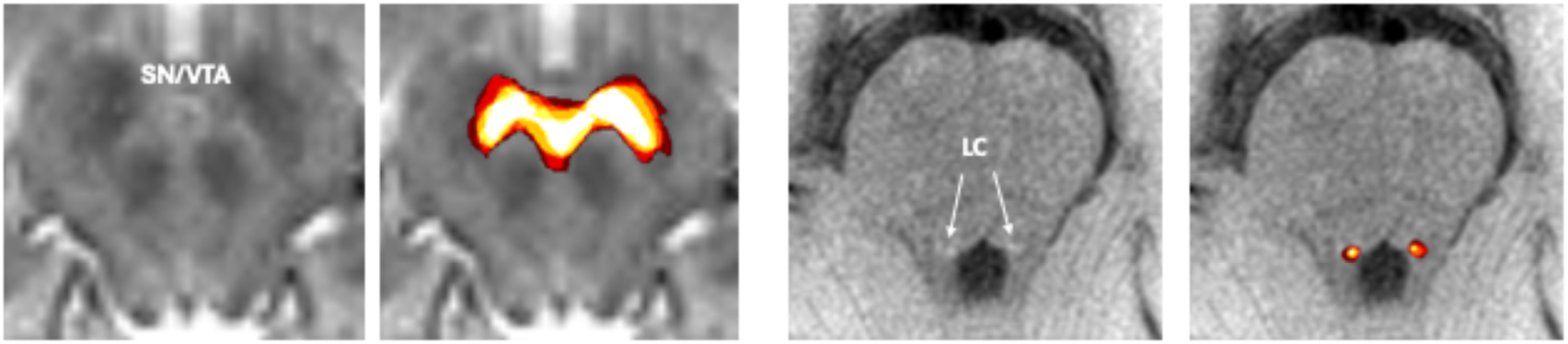
Illustration of the SN/VTA (left) and LC (right) in a representative individual and as an average mask. Note that these masks are shown for illustrative purposes only (average of n = 6 participants), as they were drawn in each participant’s native space and are therefore not directly comparable across subjects in a quantitative sense.

The average tSNR across all subjects was *M* = 33.92 (*SD* = 4.57), which is above the standard cutoff of < 30 (Drescher et al., 2026; Grueschow et al., 2021). Greenhouse-Geisser correction to degrees of freedom and *p* values was applied in case of sphericity violations. Values exceeding three standard deviations from the mean were treated as outliers and excluded from analyses. Significant effects and interactions were followed by post hoc t-tests. The statistical analyses were performed in Rstudio (R version 4.5.1).

#### 2.2.3. General Psychophysiological Interactions (gPPI)

To examine task-dependent changes in functional connectivity of the LC and SN/VTA with the rest of the brain, generalized psychophysiological interaction (gPPI) analyses were conducted using the gPPI toolbox (version 13.1; (McLaren et al., 2012) implemented in SPM12. Unlike traditional PPI, which models a single psychological contrast, gPPI simultaneously models the interaction between the seed region time series and all task conditions, thereby providing more accurate and condition-specific connectivity estimates (McLaren et al., 2012)

For each participant, two separate gPPI models were estimated, one for the LC seed and one for the SN/VTA seed, using the individually drawn masks described in section 2.2.2. The gPPI model included PPI regressors for each stimulus (reward, neutral, salience, and standard), alongside the main effects of task conditions and the physiological regressor (seed time series). The six motion parameters from the realignment procedure were included as regressors of no interest. Only PPI contrasts were defined at the first level for the significant findings of the ROI analyses. The contrast images were normalized to MNI space and subsequently spatially smoothed with a 6 mm FWHM Gaussian kernel to improve signal-to-noise ratio and inter-subject alignment for group-level inference. Note that this smoothing was applied only to the gPPI contrast images (i.e., the target side of the connectivity analysis), not to the functional data used for seed time series extraction, thereby preserving the spatial specificity of the brainstem seed regions.

At the second level, one-sample *t*-tests were performed for each PPI contrast that showed a significant effect in the preceding ROI-to-ROI analyses (see Results, section 3.2.1) separately, testing whether the group mean differed significantly from zero. This was done independently for the LC and SN/VTA seeds. Statistical maps were thresholded at *p* < .001 (uncorrected) at the voxel level with a cluster-extent threshold determined by SPM’s random field theory for FWE correction at the cluster level (*p* < .05).

## 3. Results

### 3.1 Behavioural results

Descriptive statistics for reaction times and accuracy of responses are displayed in Figure 3. Behavioural results of two participants were not recorded due to technical issues. A main effect was found for accuracy (F(2, 88) = 4.286, *p* = .017, η²_p_ = .089). Post hoc paired t-tests (Holm-Bonferroni corrected) revealed higher accuracy for rewarded oddballs (M = 85.3%, SD = 10.7) than for neutral oddballs (M = 80.3%, SD = 13.2; t(42) = 2.96, p = .015, d = 0.45). Accuracy for salient oddballs (M = 84.4%, SD = 8.8) did not differ from either rewarded (t(42) = 0.55, p = .587, d = 0.08) or neutral oddballs (t(42) = 1.95, p = .115, d = 0.30).

**Figure 3.**
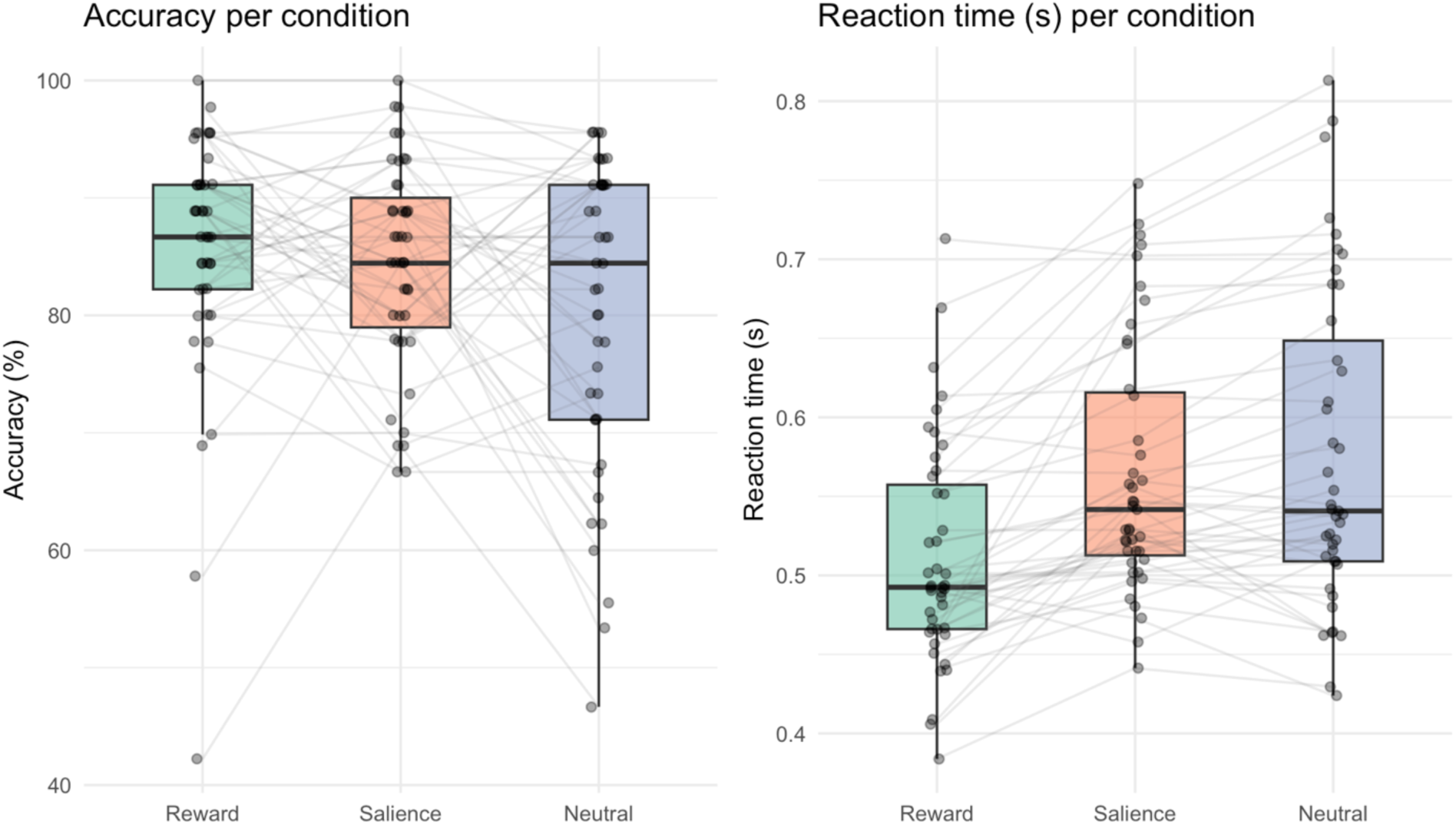
Mean accuracy (left) and reaction time (right) for each condition (oddball). Note: salience = visual salience.

Reaction times also showed a main effect of condition (F(1.59, 66.98) = 36.78, p < .001, η²p = .467, Greenhouse-Geisser corrected). Responses to rewarded oddballs (M = 512 ms, SD = 70) were faster than to neutral oddballs (M = 576 ms, SD = 101; t(42) = −6.78, p < .001, d = −1.03) and to salient oddballs (M = 565 ms, SD = 80; t(42) = −6.44, p < .001, d = −0.98). Reaction times for salient and neutral oddballs did not differ significantly (t(42) = −1.85, p = .071, d = −0.28).

### 3.2 fMRI results

#### 3.2.1 Region of interest results

Mean datapoints of the LC and SN/VTA beta values are illustrated in Figure 4. For the SN/VTA analysis, six datapoints (1.5%) were identified as outliers and were removed from the analysis. Three LC data points (0.8%) were excluded as outliers.

**Figure 4.**
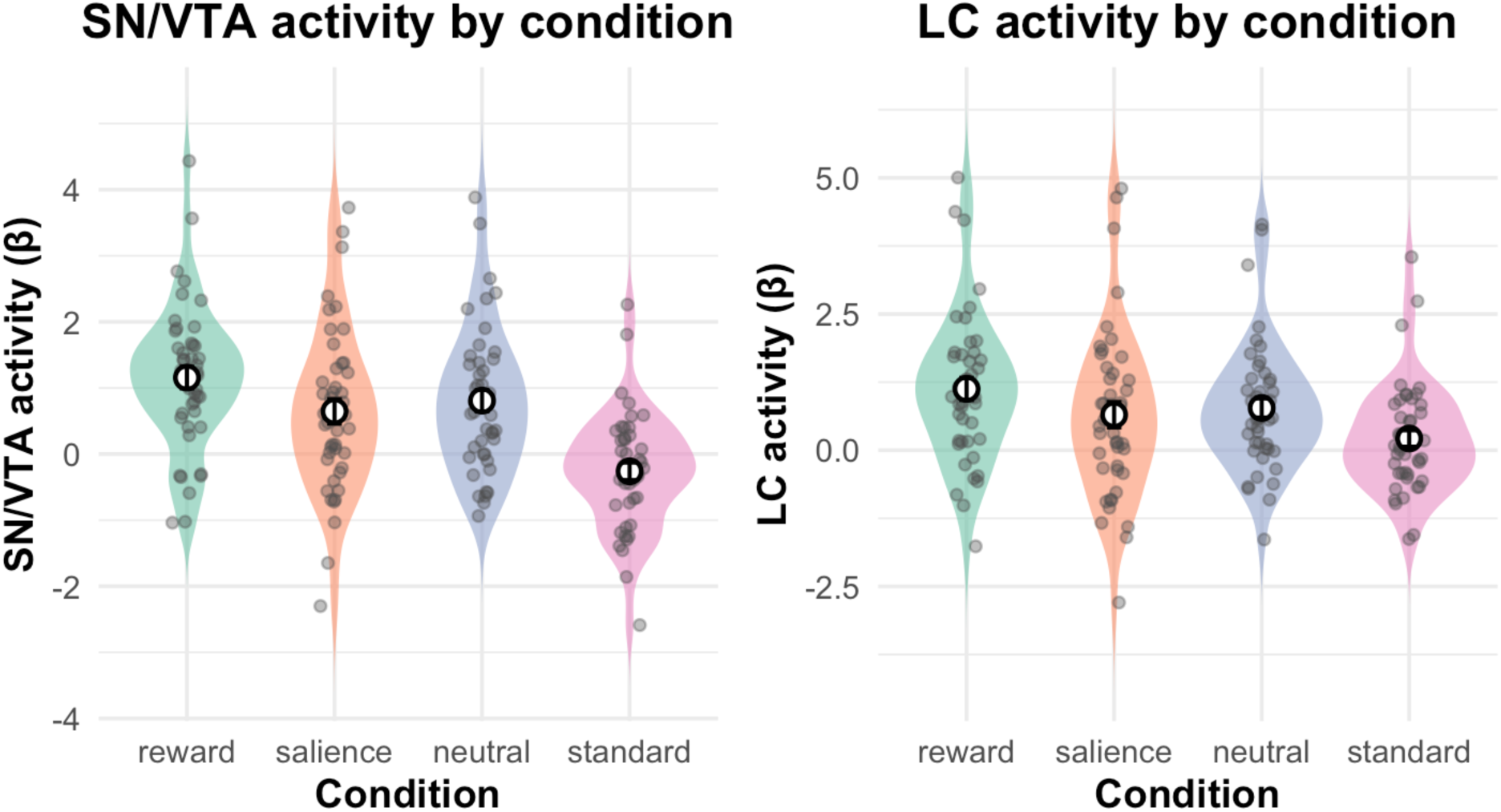
Mean beta values for each condition (oddball) of the SN/VTA (left) and the LC (right). Note: salience = visual salience.

The SN/VTA analysis revealed a main effect of stimulus type (F(2, 88) = 7.22, p = .001, η²p = .141). Post hoc paired t-tests (Holm-Bonferroni corrected) showed that SN/VTA activity was higher for rewarded oddballs (M = 1.15, SD = 1.10) than for neutral oddballs (M = 0.76, SD = 1.06; t(44) = 2.95, p = .010, d = 0.44) and salient oddballs (M = 0.64, SD = 1.25; t(44) = 3.89, p = .001, d = 0.58). Salient and neutral oddballs did not differ (t(44) = −0.81, p = .422, d = −0.12).

LC activity also showed a main effect of stimulus type (F(2, 88) = 4.21, p = .018, η²p = .087). LC activity was higher for rewarded oddballs (M = 1.13, SD = 1.37) than for salient oddballs (M = 0.65, SD = 1.55; t(44) = 2.56, p = .042, d = 0.38). The difference between rewarded and neutral oddballs (M = 0.78, SD = 1.18) did not reach significance (t(44) = 2.20, p = .066, d = 0.33), nor did the difference between salient and neutral oddballs (t(44) = −0.76, p = .449, d = −0.11).

In secondary analyses including the standard condition, both LC and SN/VTA activity was higher for all oddball types than for standards (LC standard: M = 0.22, SD = 1.01; SN/VTA standard: M = −0.26, SD = 0.88; all p ≤ .042, Holm-Bonferroni corrected).

#### 3.2.2. Generalized Psychophysiological Interactions (gPPI)

To examine whether the observed differences in LC and SN/VTA activation were accompanied by condition-dependent changes in functional connectivity, gPPI analyses were conducted for contrasts that showed significant effects in the ROI analysis. Specifically, we tested reward *vs* salience connectivity for the LC seed, and both reward *vs* neutral and reward *vs* salience connectivity for the SN/VTA seed. For the LC seed, the reward *vs* salience contrast did not reveal any significant clusters at the whole-brain level (voxel-level threshold *p* < .001, uncorrected; cluster-level *p* FWE < .05). Similarly, for the SN/VTA seed, neither the reward versus neutral nor the reward versus salience contrast yielded significant clusters at the corrected threshold.

## Discussion

While the dopaminergic system (including the SN/VTA complex) has traditionally been implicated in reward processing, and the noradrenergic system (with the LC as its main source) most prominently with processing salient information, more recent studies in both animals and humans (using fMRI and pupillometry) have shown that there is substantial overlap between these neuromodulator systems in regulating these processes and associated responses. Both systems respond to motivationally relevant as well as to more basic forms of salience, including novelty (for reviews, see Bromberg-Martin et al., 2010; Poe et al., 2020). However, in many of these studies, salience and reward are either studied in isolation or are inherently conflated, making it difficult to determine whether these systems encode similar or distinct aspects of motivational relevance. In addition, salience is used in different types of contexts, and is manipulated using different types of stimuli. With the current study we sought to systematically explore the contributions of the two neuromodulatory systems to different facets of salience processing in humans within the same paradigm. We employed three types of oddball stimuli: 1) rewarding oddballs, 2) visually salient oddballs, and 3) neutral (non-rewarding and visually non-salient) oddballs. Importantly, all oddballs were similar in terms of their predictability (i.e., equally infrequent relative to the standard), as well as their response requirements (in that all required a manual response). By comparing oddballs associated with reward prospect to visually salient oddballs without reward prospect, the present study provides insight into how reward value shapes LC–NA and SN/VTA-DA engagement above and beyond more basic forms of salience.

Reward prospect improved both accuracy and response speed relative to neutral oddballs. Visually salient oddballs, by contrast, did not reliably differ from neutral oddballs on either measure, with accuracy falling between the neutral and rewarded conditions. This pattern aligns with the well-known performance-facilitating effects of reward prospect, which are thought to reflect enhanced motivational engagement and response vigour (Braver et al., 2014). Salience alone may capture attention without providing this motivational drive.

On the neural level, we found a significant activity increase in the SN/VTA complex in the rewarded compared to both the visually salient and neutral oddball conditions. This resonates with the well-established notion of increased dopaminergic activity in response to a motivationally relevant stimulus (Braver et al., 2014; Hall et al., 2024; Krebs et al., 2012; Schultz, 1998). Moreover, we also found enhanced LC activity in the rewarded compared to the visually salient oddball condition. Reward-related LC activity was also numerically higher than in the neutral condition, though this difference did not reach significance (*p* = .066). Together, these findings suggest that reward amplifies noradrenergic activity beyond the effects of visual salience alone, potentially reflecting heightened arousal and attentional readiness when salient events carry motivational significance. Importantly, these modulations occurred in addition to a basic sensitivity of both systems to deviation from expectancy in combination with response relevance: all oddball stimuli elicited enhanced activity in both LC and SN/VTA systems compared to the standard stimuli.

An interesting dissociation emerged between behavioural performance and neuromodulatory activity. Visually salient oddballs were associated with relatively high accuracy but elicited the slowest reaction times and the (numerically) lowest activity in both LC and SN/VTA. One possible explanation is that these stimuli were perceptually easy to discriminate from other trial types, supporting accurate responses, while lacking motivational relevance to facilitate rapid responding. In contrast, neutral (no-reward) oddballs shared key stimulus features with reward trials, potentially increasing their motivational ambiguity and contributing to reduced accuracy without engaging neuromodulatory systems to the same extent as rewarded stimuli. Reaction times may therefore reflect the integration of perceptual salience and motivational relevance, with fastest responses observed when both dimensions converge, which is the case in the reward condition. This pattern is consistent with adaptive gain accounts of LC-NA function, which propose that phasic noradrenergic facilitation of behaviour is driven primarily by the decision-relevant utility of a stimulus rather than by perceptual salience alone (Aston-Jones & Cohen, 2005). This principle seems to extend to the SN/VTA-DA system given its similarly reward-selective response pattern observed here.

Whole-brain analyses (Supplementary Materials Section B) corroborated the dissociation between the reward and visually salient oddball processing. Rewarded oddball stimuli recruited a broad network of salience, affective, and cognitive-control regions (e.g., insula, mid-cingulate, amygdala, fronto-parietal cortex) beyond what was found for visually salient oddballs alone.

This confirms that reward-related oddballs are, as intended, more engaging and motivationally relevant, even if response requirements are kept equal across all oddball conditions. Notably, neutral oddballs elicited stronger activation than visually salient oddballs in several salience- and control-related regions (thalamus, pallidum, insula, mid-cingulate), which might be due to a resemblance of reward stimuli in basic shape and color which might lead to additional attentional capture and subsequent cognitive processing to reconsider the meaning. This is a post hoc interpretation that needs further exploration.

The gPPI analyses did not reveal significant condition-dependent changes in whole-brain functional connectivity for either the LC or SN/VTA seed regions. Given the widespread cortical projections of both nuclei, it is unlikely that these regions operate in isolation during reward and salience processing. Rather, the absence of significant results likely reflects a limitation in detection power. (g)PPI analyses are known to have substantially lower statistical power compared to standard activation analyses, as the PPI regressor represents an interaction term that captures only a small proportion of the total variance in the BOLD signal. Although our power analysis was adequate for the performance and ROI analysis, the event-related oddball design included a relatively low number of trials per condition (45 per condition across three blocks), which may have been insufficient to reliably estimate condition-specific connectivity changes.

For comparison, we previously employed 72 trials per condition and observed significant LC connectivity modulations with occipital cortex using the same gPPI approach (Hall et al., 2024). The substantially lower trial count in the current design likely reduced our ability to detect similar effects (Cisler et al., 2014; McLaren et al., 2012), suggesting that a larger number of trials per participant may have been required to achieve adequate power for the connectivity analysis. Although we were unable to address this question with the current data, it remains a compelling avenue for future research. Additionally, the anatomical properties of SN/VTA and LC, which are small and in the vicinity of blood vessels, may have contributed to a noisier seed time series (see Hall et al., 2024, for a detailed discussion). This is a general concern for studying small regions in the vicinity of blood vessels brainstem. However, the tSNR was above the cut-off score (Grueschow et al., 2021). Studies with higher trial counts and potentially block designs, which afford greater power for connectivity analyses, might be more sensitive for functional connectivity of LC and SN/VTA in the context of salience processing.

Further, the fixed visual identity of the oddball stimuli might present a design limitation: because all oddballs were consistently represented by the same shape and colour (e.g., a grey triangle for the rewarded oddball) across the task, participants may have habituated to these repeated low-level features over time, potentially attenuating condition-specific salience or novelty effects, particularly for the visually salient oddball. Varying the colour or form of the oddballs across trials, while holding condition-defining features constant, may have yielded even greater experimental control of perceptual salience throughout the task. Relatedly, because the reward and neutral oddballs were similar in basic shape and colour (differing only in triangle orientation), participants had to rely on the orientation, perhaps a relatively subtle featural distinction, to differentiate these conditions, which may have introduced additional discrimination demands or transient ambiguity. While this design choice helped minimise low-level visual differences between reward and neutral trials, it may also have made the two conditions harder to distinguish than the visually salient oddball, which differed from the standard more categorically (i.e., by colour).

## Conclusion

Taken together, this study shows that the LC-NA and SN/VTA-DA complex are both sensitive to motivationally relevant stimuli, but do not respond to reward and salience in an identical manner. Behaviourally, reward prospect facilitates goal-directed responding beyond what perceptual salience alone can achieve, reflecting typical instrumental response invigoration. This was mirrored at the neural level: LC activity was enhanced for rewarded compared to visually salient oddballs, and SN/VTA activity was likewise higher for rewarded compared to both neutral and visually salient conditions. At the same time, the dissociation observed between neutral and visually salient oddballs, which were associated with high accuracy but slower responses and comparatively low LC and SN/VTA activity, and neutral oddballs suggests that perceptual ease and motivational relevance contribute independently to behaviour, with the fastest responses emerging only when both dimensions converge, i.e., in the reward condition. These findings extend prior work on reward and salience processing by demonstrating that both the noradrenergic and dopaminergic system jointly contribute to goal-directed responding to motivationally relevant stimuli – exceeding the processing of more basic forms of salience. Future work using designs that vary stimulus identity more flexibly across trials, and that are adequately powered for connectivity analyses, will be needed to clarify the extent to which the LC and SN/VTA operate through shared pathways in the service of adaptive, reward-guided action.

## Data and code availability

Data is available after submission of a formal (brief) project outline to the corresponding author and after a formal data sharing agreement has been signed. Code is available on github.com/juliemaehall/LC_VTA_Reward_Salience

## Author statement

Julie M. Hall: Conceptualization, Methodology, Investigation, Formal Analysis, Writing - Original Draft, Visualization, Funding Acquisition.

Nanne Kukkonen: Investigation, Writing - Reviewing and Editing. Eleni Prinea: Formal Analysis, Writing-Reviewing and Editing Judith Schomaker: Writing - Reviewing and Editing

Ruth Krebs: Conceptualization, Supervision, Writing - Reviewing and Editing, Funding Acquisition

## Acknowledgements

We thank Fearghal Lee for his contribution to the ROI delineation, and all volunteer participants in this study.

