## Supplementary Materials for "The Role of Locus Coeruleus and Substantia Nigra/Ventral Tegmental Area activity in Reward and Salience Processing: A High-Resolution fMRI Study"

**Section A. Automated boilerplate text output, excluding information not pertaining to the output used in the present study**

Results included in this manuscript come from preprocessing performed using *fMRIPrep* 21.0.1 (Esteban et al., 2019); RRID:SCR_016216), which is based on *Nipype* 1.6.1 (Gorgolewski et al., 2011); (Gorgolewski et al., 2018), RRID:SCR_002502).

**Preprocessing of B_0_ inhomogeneity mappings**
A *B_0_* nonuniformity map (or *fieldmap*) was estimated from the phase-drift map(s) measure with two consecutive GRE (gradient-recalled echo) acquisitions. The corresponding phase-map(s) were phase-unwrapped with prelude (FSL 6.0.5.1:57b01774).

**Anatomical data preprocessing**

The T1-weighted (T1w) images were corrected for intensity non-uniformity (INU) with N4BiasFieldCorrection (Tustison et al., 2010), distributed with ANTs 2.3.3 [(Avants et al., 2008), RRID:SCR_004757)], and used as T1w-reference throughout the workflow. The T1w-reference was then skull-stripped with a *Nipype* implementation of the antsBrainExtraction.sh workflow (from ANTs), using OASIS30ANTs as target template. Brain tissue segmentation of cerebrospinal fluid (CSF), white-matter (WM) and gray-matter (GM) was performed on the brain-extracted T1w using fast (FSL 6.0.5.1:57b01774, RRID:SCR_002823) (Zhang et al., 2001). Volume-based spatial normalization to two standard spaces (MNI152NLin2009cAsym, MNI152NLin6Asym) was performed through nonlinear registration with antsRegistration (ANTs 2.3.3), using brain-extracted versions of both T1w reference and the T1w template. The following templates were selected for spatial normalization: *ICBM 152 Nonlinear Asymmetrical template version 2009c*[(Fonov et al., 2009), RRID:SCR_008796; TemplateFlow ID: MNI152NLin2009cAsym], *FSL’s MNI ICBM 152 non-linear 6th Generation Asymmetric Average Brain Stereotaxic Registration Model* [Evans et al. (2012), RRID:SCR_002823; TemplateFlow ID: MNI152NLin6Asym].

**Section B. Whole brain analysis and results
Functional data preprocessing**

For each of the 3 BOLD runs found per subject (across all tasks and sessions), the following preprocessing was performed. First, a reference volume and its skull-stripped version were generated using a custom methodology of *fMRIPrep*. Head-motion parameters with respect to the BOLD reference (transformation matrices, and six corresponding rotation and translation parameters) are estimated before any spatiotemporal filtering using mcflirt (FSL 6.0.5.1:57b01774 (Jenkinson et al., 2002)). BOLD runs were slice-time corrected to 0.99s (0.5 of slice acquisition range 0s-1.98s) using 3dTshift from AFNI (Cox & Hyde, 1997), RRID:SCR_005927). The BOLD time-series (including slice-timing correction when applied) were resampled onto their original, native space by applying the transforms to correct for head-motion. These resampled BOLD time-series will be referred to as *preprocessed BOLD in original space*, or just *preprocessed BOLD*. The BOLD reference was then co-registered to the T1w reference using mri_coreg (FreeSurfer) followed by flirt (FSL 6.0.5.1:57b01774, Jenkinson & Smith, 2001)) with the boundary-based registration (Greve & Fischl, 2009) cost-function. Co-registration was configured with six degrees of freedom. Several confounding time-series were calculated based on the *preprocessed BOLD*: framewise displacement (FD), DVARS and three region-wise global signals. FD was computed using two formulations following Power (absolute sum of relative motions, Power et al. (2014) and Jenkinson (relative root mean square displacement between affines (Jenkinson et al., 2002). FD and DVARS are calculated for each functional run, both using their implementations in *Nipype* (following the definitions by Power et al., 2014). The three global signals are extracted within the CSF, the WM, and the whole-brain masks. Additionally, a set of physiological regressors were extracted to allow for component-based noise correction (*CompCor*, Behzadi et al., 2007). Principal components are estimated after high-pass filtering the *preprocessed BOLD* time-series (using a discrete cosine filter with 128s cut-off) for the two *CompCor* variants: temporal (tCompCor) and anatomical (aCompCor). tCompCor components are then calculated from the top 2% variable voxels within the brain mask. For aCompCor, three probabilistic masks (CSF, WM and combined CSF+WM) are generated in anatomical space. The implementation differs from that of Behzadi et al. in that instead of eroding the masks by 2 pixels on BOLD space, the aCompCor masks are subtracted a mask of pixels that likely contain a volume fraction of GM. This mask is obtained by thresholding the corresponding partial volume map at 0.05, and it ensures components are not extracted from voxels containing a minimal fraction of GM. Finally, these masks are resampled into BOLD space and binarized by thresholding at 0.99 (as in the original implementation). Components are also calculated separately within the WM and CSF masks. For each CompCor decomposition, the *k* components with the largest singular values are retained, such that the retained components’ time series are sufficient to explain 50 percent of variance across the nuisance mask (CSF, WM, combined, or temporal). The remaining components are dropped from consideration. The head-motion estimates calculated in the correction step were also placed within the corresponding confounds file. The confound time series derived from head motion estimates and global signals were expanded with the inclusion of temporal derivatives and quadratic terms for each (Satterthwaite et al., 2013). Frames that exceeded a threshold of 0.5 mm FD or 1.5 standardised DVARS were annotated as motion outliers. For the ROI analysis, images were spatially smoothed with Gaussian an kernel of 3.5 mm FWHM (full-width half-maximum). For the whole brain analysis, automatic removal of motion artifacts using independent component analysis (ICA-AROMA, Pruim et al. 2015) was performed on the *preprocessed BOLD on MNI space* time-series after removal of non-steady state volumes and spatial smoothing with an isotropic, Gaussian kernel of 6mm FWHM. Corresponding “non-aggresively” denoised runs were produced after such smoothing. Additionally, the “aggressive” noise-regressors were collected and placed in the corresponding confounds file. All resamplings can be performed with *a single interpolation step* by composing all the pertinent transformations (i.e. head-motion transform matrices, susceptibility distortion correction when available, and co-registrations to anatomical and output spaces). Gridded (volumetric) resamplings were performed using antsApplyTransforms (ANTs), configured with Lanczos interpolation to minimize the smoothing effects of other kernels (Lanczos 1964). Non-gridded (surface) resamplings were performed using mri_vol2surf (FreeSurfer).

**Methods whole brain analysis**

***Second level analysis***

Voxel-wise random-effects analyses were performed for the reward oddball paradigm using a flexible factorial design, with stimulus type (reward/neutral/visual salience) as factor. We compared reward vs neutral, reward vs visual salience and neutral vs visual salience. Activations were considered significant if they survived a cluster-level family-wise error (FWE) correction of *p* <. 05, based on an uncorrected voxel-wise threshold of *p* < .001.

**Whole brain results**

*Reward vs. neutral oddball*

For the reward *vs.* neutral contrast in the oddball task, whole-brain analyses revealed widespread activation encompassing fronto-cingulate regions, bilateral temporal and occipital cortices, and subcortical structures including the amygdala, thalamus, caudate, and pallidum (all *p* < .05 FWE-corrected). Effects were also observed in limbic (parahippocampal, fusiform, insula), parietal (angular gyrus, precuneus), sensorimotor, brainstem/midbrain, and cerebellar regions, indicating broad engagement of reward, salience, and perceptual networks during rewarded oddball processing. Table S1 provides a detailed overview of the significant regions in this contrast.

**Table S1. Whole brain results in the reward *vs* neutral contrast**

| **Region** | **x** | **y** | **z** | **k** | **t-value** | ***p* FWE** |
| --- | --- | --- | --- | --- | --- | --- |
| R Mid Cingulum | 4 | -32 | 30 | 2101 | 8.04 | <.001 |
| L Angular | -56 | -60 | 28 | 1459 | 6.91 | <.001 |
| R Mid Temporal | 56 | -18 | -10 | 2017 | 6.62 | <.001 |
| L Mid Temporal | -66 | -28 | -8 | 844 | 6.51 | <.001 |
| R Amygdala | 26 | -2 | -16 | 604 | 6.44 | <.001 |
| L Fusiform | -34 | -54 | -16 | 46 | 6.26 | .031 |
| L Insula | -28 | 20 | -4 | 74 | 6.19 | .002 |
| L Mid Occipical | -20 | -94 | 12 | 329 | 5.97 | <.001 |
| L Superior Occipital | -24 | -80 | 32 | 318 | 5.49 | <.001 |
| L Caudate | -10 | 10 | -4 | 95 | 5.47 | <.001 |
| R Thalamus | 4 | -6 | 4 | 271 | 5.37 | <.001 |
| L Parahippocampal | -24 | -26 | -14 | 137 | 5.37 | <.001 |
| R Fusiform | 26 | -48 | -16 | 950 | 5.27 | <.001 |
| L Superior Frontal | -12 | 34 | 54 | 653 | 5.22 | <.001 |
| Midbrain | 6 | -18 | -18 | 132 | 5.22 | <.001 |
| R Cerebelum Crus2 | 20 | -28 | 58 | 50 | 5.21 | .02 |
| L Lingual | -28 | -86 | -16 | 465 | 5.14 | <.001 |
| R Cerebelum Crus2 | 20 | -82 | -34 | 231 | 5.13 | <.001 |
| L Cerebelum 9 | -8 | -48 | -42 | 177 | 5.06 | <.001 |
| L Inf Frontal Trt | -52 | 24 | 2 | 71 | 5.02 | .003 |
| L Inf Fronal Orbital | -46 | 44 | -14 | 75 | 5.00 | .002 |
| L Precuneus | -6 | -70 | 34 | 53 | 5.00 | .015 |
| L Superior Frontal | -20 | 56 | 30 | 69 | 4.98 | .003 |
| L Cerebelum 4,5 | -14 | -36 | -20 | 133 | 4.95 | <.001 |
| R Cerebelum 4,5 | 26 | -40 | -30 | 162 | 4.92 | <.001 |
| Brainstem | 2 | -26 | -44 | 46 | 4.88 | .031 |
| Vermis 7 | -2 | -62 | -24 | 46 | 4.82 | .031 |
| L Mid Frontal | -48 | 16 | 46 | 92 | 4.79 | <.001 |
| R Mid Frontal | 46 | 14 | 50 | 96 | 4.66 | <.001 |
| L Pallidum | -24 | -2 | 4 | 55 | 4.66 | .012 |
| L Mid Frontal Orbital | -22 | 32 | -14 | 42 | 4.66 | .047 |
| L Postcentral | -40 | -16 | 32 | 76 | 4.62 | .002 |
| L Cerebelum Crus2 | -36 | -68 | -42 | 46 | 4.52 | .031 |
| R Postcentral | 48 | -8 | 30 | 78 | 4.45 | .001 |

*Notes. H = hemisphere (L = left, R = right); x;y;z = coordinates in MNI space. k = cluster size. Brain regions labels represent local maxima. Only clusters surviving an FWE-corrected threshold of p < .05 and consisting of at least five voxels (p < .001) are shown.*

*Reward vs. visual salience oddball*

Table S2 provides a detailed overview of the significant regions in the reward *vs.* (visual) visual salience contrast. Here we found activation in salience and affective regions, including the right insula, mid-cingulate cortex, and amygdala, alongside engagement of fronto-parietal control areas (middle frontal and inferior parietal cortices). Significant effects were also detected in occipito-temporal visual regions, sensorimotor cortex, and subcortical structures such as the putamen, reflecting a combination of **perceptual, motivational, and action-related processing.** These results indicate that reward-related oddballs recruit widespread networks beyond visual salience, encompassing motivational, affective, and cognitive control systems (all p < .05 FWE-corrected).

**Table S2. Whole brain results in the reward *vs* visual salience contrast**

| **Region** | **x** | **y** | **z** | **k** | **t-value** | ***p* FWE** |
| --- | --- | --- | --- | --- | --- | --- |
| R Insula | 40 | 18 | -6 | 695 | 8.22 | <.001 |
| R Mid Cingulum | 2 | -20 | 32 | 5198 | 8.14 | <.001 |
| L Amygdala | -14 | 6 | -8 | 7847 | 7.57 | <.001 |
| R Inf Occipital | 38 | -84 | 0 | 3161 | 6.14 | <.001 |
| R Inf Parietal | 52 | -44 | 40 | 1377 | 5.90 | <.001 |
| L Mid Temporal | -52 | -30 | -6 | 228 | 5.27 | <.001 |
| L Cerebelum 4,5 | -8 | -44 | -18 | 91 | 5.12 | <.001 |
| R Sup Temporal | 50 | -26 | -8 | 425 | 5.06 | <.001 |
| L Inf Parietal | -56 | -50 | 38 | 613 | 5.04 | <.001 |
| L Postcentral | -42 | -14 | 30 | 145 | 4.96 | <.001 |
| R Mid Frontal | 48 | 18 | 42 | 528 | 4.87 | <.001 |
| R Postcentral | 30 | -22 | 46 | 154 | 4.68 | <.001 |
| R Postcentral | 46 | -16 | 36 | 110 | 4.68 | <.001 |
| L Putamen | -32 | -16 | 0 | 93 | 4.63 | <.001 |
| L Mid Frontal | -36 | 26 | 44 | 136 | 4.31 | <.001 |
| L Mid Temporal | -62 | -20 | -16 | 42 | 4.24 | .047 |
| L Inf Frontal Oper | -56 | 12 | 14 | 46 | 4.11 | .031 |

*Notes. H = hemisphere (L = left, R = right); x;y;z = coordinates in MNI space. k = cluster size. Brain regions labels represent local maxima. Only clusters surviving an FWE-corrected threshold of p < .05 and consisting of at least five voxels (p < .001) are shown.*

*Neutral vs visual salience oddball*

For the neutral oddball > visual salience contrast, whole-brain analyses showed significant activation in the bilateral thalamus, pallidum, bilateral insula, and mid-cingulate cortex, as well as brainstem and cerebellar Crus I (all *p* < .05 FWE-corrected). Although the visually salient stimuli were designed to maximize salience, neutral oddballs elicited stronger activation in salience- and control-related regions, likely reflecting their greater behavioural relevance as they more closely resemble the reward oddball, rather than low-level visual salience per se.

**Table S3. Whole brain results in the neutral vs perceptual salience contrast**

| **Region** | **x** | **y** | **z** | **k** | **t-value** | ***p* FWE** |
| --- | --- | --- | --- | --- | --- | --- |
| R Thalamus | 14 | -12 | 2 | 329 | 5.45 | <.001 |
| L Pallidum | -16 | 4 | 4 | 136 | 5.37 | <.001 |
| Brainstem | -8 | -22 | -28 | 134 | 5.06 | <.001 |
| Cerebellum crus 1 | -34 | -54 | -36 | 152 | 5.02 | <.001 |
| R Mid Cingulum | 2 | 10 | 46 | 247 | 4.99 | <.001 |
| R Insula | 36 | 16 | -4 | 48 | 4.87 | .025 |
| L Insula | -38 | 14 | 8 | 119 | 4.51 | <.001 |
| L Thalamus | -14 | -20 | 10 | 43 | 4.27 | .042 |

*Notes. H = hemisphere (L = left, R = right); x;y;z = coordinates in MNI space . k = cluster size. Brain regions labels represent local maxima. Only clusters surviving an FWE-corrected threshold of p < .05 and consisting of at least five voxels (p < .001) are shown.*
